# Microevolutionary cophylogeny reflects host-symbiont population dynamics and human mitonuclear interactions

**DOI:** 10.64898/2026.09.11.751054

**Authors:** Rowan Hart, Matthias Steinrücken

## Abstract

Cophylogeny, the study of phylogenetic similarity between interacting organisms, provides insights into the specificity and shared evolutionary history of symbiosis. While the ecological drivers of cophylogeny have been investigated at the macroevolutionary scale, the influence of these processes on microevolution remains unclear. This is due, in part, to the fact that the ancestral relations between individuals within a sexually reproducing eukaryotic host species cannot be well represented with a single phylogenetic tree, since genetic distances between individuals change substantially across the genome due to meiotic recombination. This heterogeneity can be captured and utilized through the inference of an ancestral recombination graph (ARG) built from the genomic data of the host. Here, we propose to measure microevolutionary cophylogeny by comparing a symbiont evolutionary tree to a host ARG. This approach simultaneously measures genome-wide cophylogeny, as well as locus-specific signals. Through simulations, we investigate the effects of transmission mode, population structure, admixture, and allelic incompatibility on microevolutionary cophylogeny. In contrast to macroevolutionary patterns, we find a limited relationship between cophylogeny and vertical transmission, with vertically transmitted host-symbiont systems displaying no cophylogeny in large panmictic populations. We apply our approach to mitochondrial and nuclear genomes within the 1000 Genomes Project— a host-symbiont system with strict maternal transmission—and observe substantial variation in mitochondrial-nuclear (mitonuclear) cophylogeny across human populations. Finally, we investigate locus-specific signals of cophylogeny and observe limited evidence of mitonuclear incompatibility.

## 1 Introduction

Cophylogeny, the measurement of phylogenetic similarity between ecologically interacting organisms, can provide insights into the evolutionary history of symbiosis and host specificity (Moran et al. 2008; Moran et al. 2019; Herńandez-Herńandez et al. 2021). Cophylogeny has primarily been measured at the macroevolutionary scale, and much work has gone into developing tools and methods designed to investigate species level processes (Vienne et al. 2013; Dismukes et al. 2022). Despite these advances, linking ecological mechanisms to cophylogenetic signals remains an outstanding challenge in cophylogenetic studies, and the connection between microevolutionary processes and macroevolutionary patterns remains to be fully understood (Blasco-Costa et al. 2021). Measuring cophylogeny at the microevolutionary scale is an important step towards understanding the relationship between ecological interactions and cophylogeny, as the mechanisms that cause cophylogeny between hosts and symbionts remain unclear in complex systems (Suzuki et al. 2022; Moeller et al. 2023; Good 2026).

In previous cophylogenetic studies, the heterogeneity of phylogenies across genomic regions has been largely underutilized (Blasco-Costa et al. 2021). This is especially important for the microevolution of sexually reproducing eukaryotes, since genealogical relations between individuals change substantially across the genome due to meiotic recombination (Wong et al. 2024). These complex genealogies can be inferred and represented with an ancestral recombination graph (ARG), which provides locus-specific genealogies across the genome for a sample of host individuals (Nielsen et al. 2025). Here, we develop an approach to measure cophylogeny between a host ARG and a symbiont evolutionary tree. This approach allows microevolutionary cophylogeny to be measured across genomic loci in the host, which separates genome-wide signals of cophylogeny from locus-specific signals. Through simulations, we explore how population structure, transmission mode, admixture, and allelic incompatibility shape cophylogeny between a symbiont tree and a host ARG.

Vertical transmission, the transfer of symbionts from parent to offspring, is a key driver of cophylogeny at the macroevolutionary scale (Moran et al. 2008; Hayward et al. 2021). Whether this association extends to microevolutionary systems has not been explored in detail. Strict vertical transmission leads to congruent pedigrees and cophylogeny between symbionts and maternally transmitted host organelles (Funk et al. 2000; Brandvain et al. 2011; Richardson et al. 2012). However, the relationship between congruent pedigrees and cophylogeny between symbionts/organelles and host nuclear loci is not fully understood (Moeller et al. 2023). Here, we demonstrate through simulations that vertical transmission does not lead to cophylogeny between symbiont trees and host nuclear loci in large panmictic populations. Rather, we find that genome-wide cophylogeny is generated through shared population structure between hosts and symbionts. We then measure cophylogeny between mitochondrial and nuclear genomes in the 1000 Genomes Project (1KGP), a genomic dataset of human samples from 26 different geographic locations (Auton et al. 2015; Byrska-Bishop et al. 2022), and observe substantial variation in mitonuclear cophylogeny across populations. Putatively unstructured populations display no genome-wide signal of cophylogeny, despite strict vertical transmission, while putatively structured populations display elevated cophylogenetic signal genome-wide.

Finally, we explore locus-specific cophylogeny and its variation across the genome. Through simulations, we demonstrate how allelic incompatibilities between hosts and symbionts can generate transient locus-specific signals at the causal locus in the host. We investigate the strongest locus-specific signals of cophylogeny within 1KGP populations and find no systematic enrichment of functional annotations. Investigating individual signals, however, reveals a potential mitonuclear interaction in Japanese in Tokyo, Japan samples at *NDUFS7*, a nuclear-encoded gene in complex I of the mitochondrial respiratory chain linked to mitochondrial diseases in humans (Lebon et al. 2007).

## 2 Results

### 2.1 Microevolutionary cophylogeny

Cophylogeny has been a powerful tool in discovering evolutionary links between host-symbiont systems across diverse taxa and evolutionary timescales. Here, we investigate microevolutionary cophylogeny by considering the entire ancestral recombination graph (ARG) of the host, and compare a single symbiont evolutionary tree to all local genealogies across the host genome encoded in the ARG (Figure 1). This approach allows for the assessment of genome-wide trends, as well as locus-specific signals in the host genome. First, we apply this approach to simulated datasets, where the host ARG and symbiont tree are known. We then apply this approach empirically to mitochondrial and nuclear genomes from the 1KGP, using `Relate` (Speidel et al. 2019) to infer ARGs from human nuclear genomes and unweighted pair group method with arithmetic mean (UPGMA; Felsenstein 2004, Ch. 11) to infer the mitochondrial tree (Section 4.2). We assess cophylogeny between the symbiont tree and the genealogies of the host ARG using `TreeDistance` (Smith 2020), a metric that quantifies topological distance between two phylogenetic trees by comparing shared bipartitions, with no consideration of branch lengths. Throughout this work, we present genome-wide cophylogeny as “cophylogeny scores” and locus-specific cophylogenetic signals as *p*-values, indicating the strength of evidence for cophylogeny; significance is assessed in both cases using permutation testing (Section 4.3).

**Figure 1:**
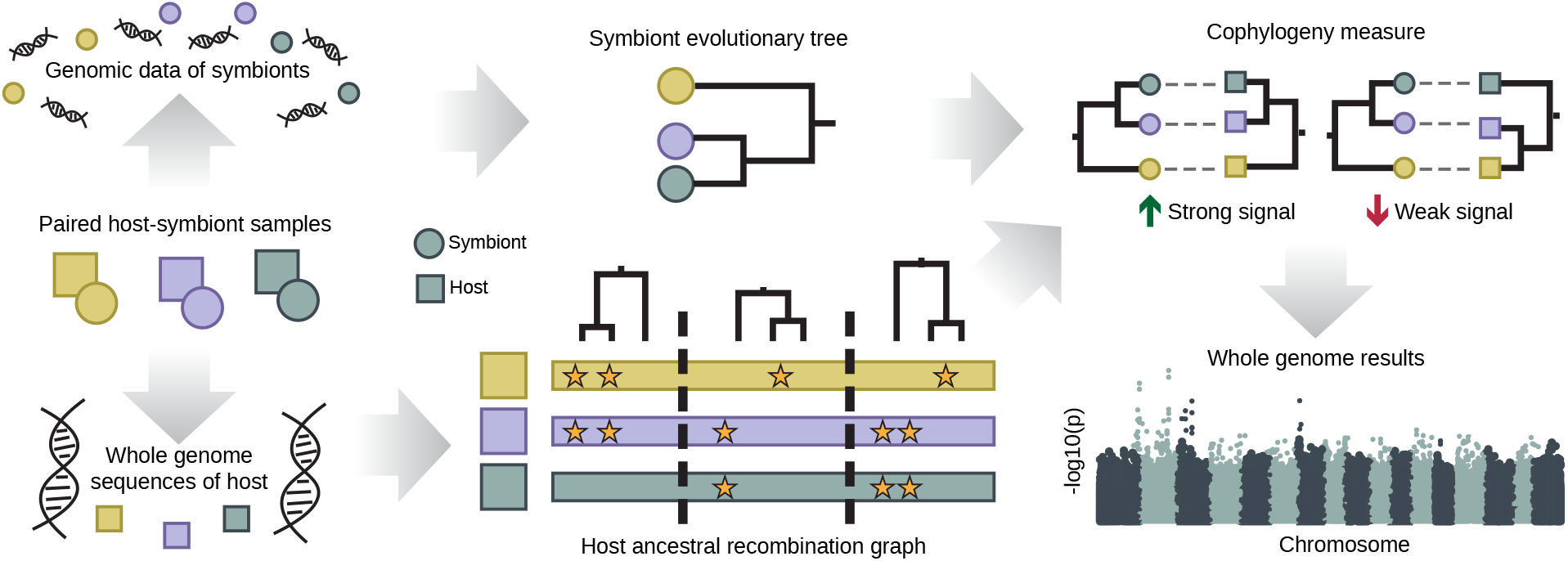
Microevolutionary cophylogeny using ancestral recombination graphs. A diagram detailing our approach to measure microevolutionary cophylogeny using an ancestral recombination graph (ARG) for the host. First, genomic data is obtained for ecologically interacting host-symbiont pairs. An evolutionary tree is built from the symbiont genomic data, and an ARG is built from the host genomic data. The strength of cophylogeny is measured between the symbiont tree and each genealogy in the host ARG, resulting in locus-specific genome-wide measurements of cophylogeny. In the bottom right panel, the genomic position of each genealogy in the host ARG is on the x-axis and the significance of cophylogeny between that genealogy and the symbiont tree is on the y-axis.

### 2.2 Vertical transmission

To explore microevolutionary cophylogeny, we developed a forward in time evolutionary model of host-symbiont evolution based on the Wright-Fisher model, a frequently used framework to model population genetic dynamics. In the model, hosts are diploid, sexual, and form discrete, non-overlapping generations with a constant population size (*N*). Host offspring select their parents by randomly sampling a male and female from the previous generation, and the sex of the host offspring is assigned at random. Each host is paired with a single symbiont. Symbionts are haploid and also form discrete, non-overlapping generations with a constant population size (*N*). Host offspring obtain their symbiont by choosing one from the previous generation, either from the mother, reflecting vertical transmission, or from a random individual, reflecting horizontal transmission (Figure 2A). Host genomes are 10 megabases (Mb) in size with a recombination rate of 10*^−^*^8^. We extract the resulting ARG for the host from each simulated replicate. Symbiont genomes are 10 megabases (Mb) in size and non-recombining. We simulated 32 replicates for each parameter combination, each of which resulted in a complete ARG for the host and a single symbiont tree. The model was implemented and run using SLiM 4.0 (Haller and Messer 2023).

**Figure 2:**
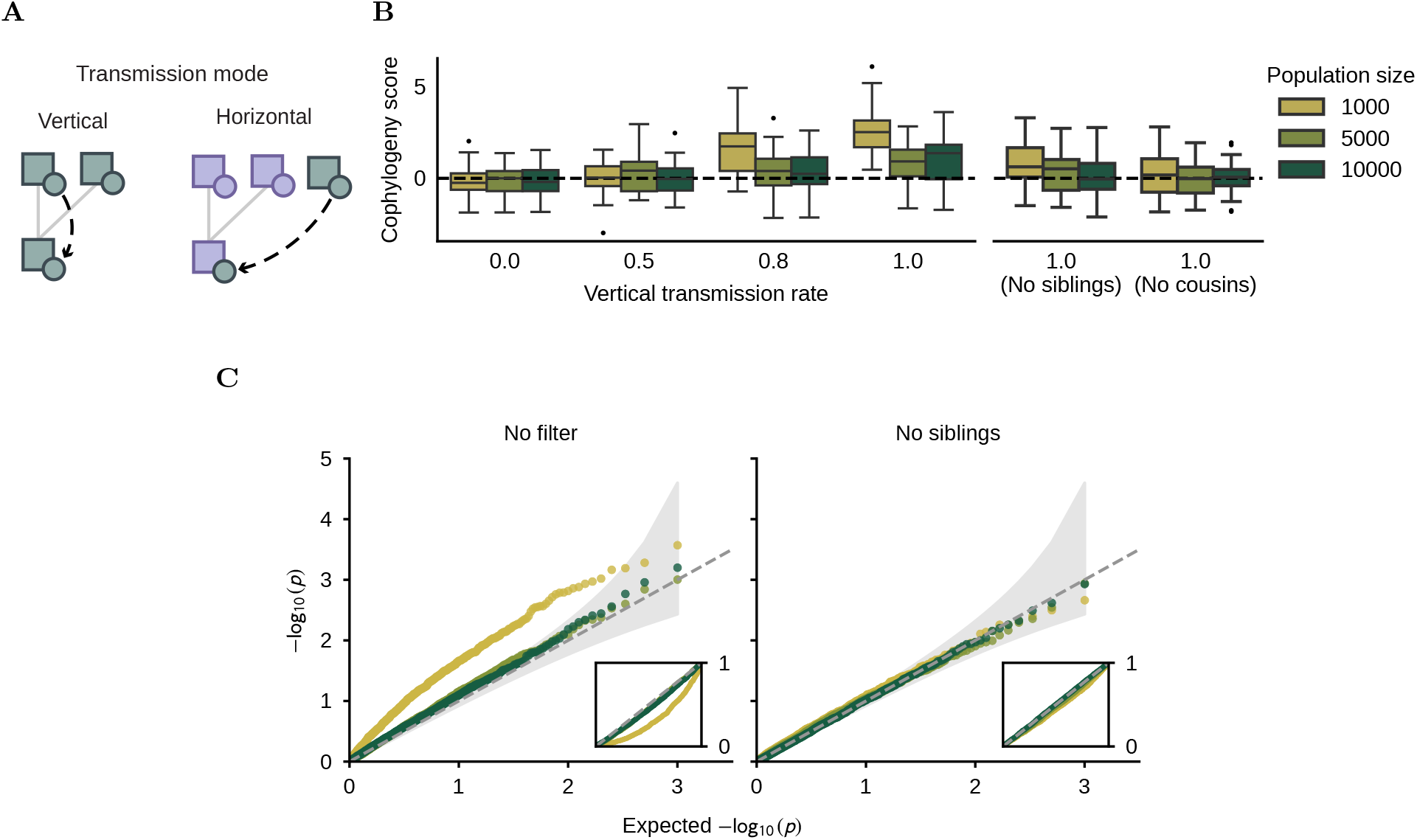
Effect of transmission mode on cophylogeny. A) Diagram of vertical and horizontal transmission in our model. The symbiont of the host offspring is either obtained from the mother, or from a different host individual. B) The effect of vertical transmission on cophylogeny scores. Each boxplot represents the cophylogeny scores of 32 simulated replicates across varying rates of vertical transmission, colored by population size. Boxplots on the right side represent analyses from simulations where individuals who shared a parent (no siblings), or individuals who shared a parent or a grandparent (no cousins) were removed. C) The effect of vertical transmission on locus-specific cophylogeny. Q-Q plots for simulations with strict vertical transmission. Cophylogenetic *p*-values were sampled at every 1 kilobase across simulated genomes and ranked by significance. Each point in the plot reflects the median ranked *p*-value across all 32 simulated replicates per parameter set. Inset Q-Q plots display the raw *p*-values. Shaded regions represent the 95% confidence intervals of expected *p*-values under the null hypothesis. Together, these plots demonstrate how vertical transmission alone does not generate cophylogeny in large panmictic populations.

We simulated an unstructured population with varying rates of vertical transmission (0.0; 0.5; 0.8; 1.0), reflecting the percentage of offspring each generation that obtain their symbiont through vertical transmission, and varying population size *N* (1, 000; 5, 000; 10, 000). Each simulation was run for 15*∗N* generations, after which we sampled 100 individuals and their symbionts, extracted the host ARGs and symbiont trees of the sampled individuals, filtered uncoalesced trees from the host ARGs (*<* 0.3%), and measured cophylogeny. For simulations with population size *N* = 10, 000, the vertical transmission rate did not affect genome-wide signals of cophylogeny between hosts and symbionts substantially, even in the case of strict vertical transmission. In small populations (*N* = 1, 000), on the other hand, we observe a slight increase of cophylogeny scores with increasing rate of vertical transmission (Figure 2B). We hypothesized that this signal arose from very recent coalescent events among closely related individuals. Indeed, when individuals who shared at least a single parent (no siblings), or individuals who shared at least a single parent or a grandparent (no cousins) were removed from our analysis, the genome-wide cophylogenetic signal vanished, even in small populations. We observed similar results for locus-specific *p*-values: inflated *p*-values in smaller populations, and no inflation after filtering for related individuals. (Figure 2C).

### 2.3 Shared population structure

To investigate the influence of population structure on cophylogeny, we extended our evolutionary model to simulate non-random mating of host individuals. To model long-standing population structure, we simulated two subpopulations with varying rates of gene flow (per generation per individual probability of switching subpopulations: 0.0005; 0.001; 0.005; 0.01) and constant population size (*N* = 5, 000 for each subpopulation) for 40 *∗ N* generations (Figure 3A). To model divergence, we simulated an ancestral host population (*N* = 10, 000) for 15 *∗ N* generations, followed by a split into two reproductively isolated subpopulations of equal size (*N* = 5, 000) (Figure 3A). In both models, the transmission mode was either strictly vertical, allopatric horizontal (symbionts horizontally transmitted within the same subpopulation), or sympatric horizontal (symbionts horizontally transmitted freely across subpopulations). To measure cophylogeny, we sampled 100 individuals and their symbionts from each subpopulation. In the population split model, sampling occurred at multiple generations following the split (1; 10; 100; 1000).

**Figure 3:**
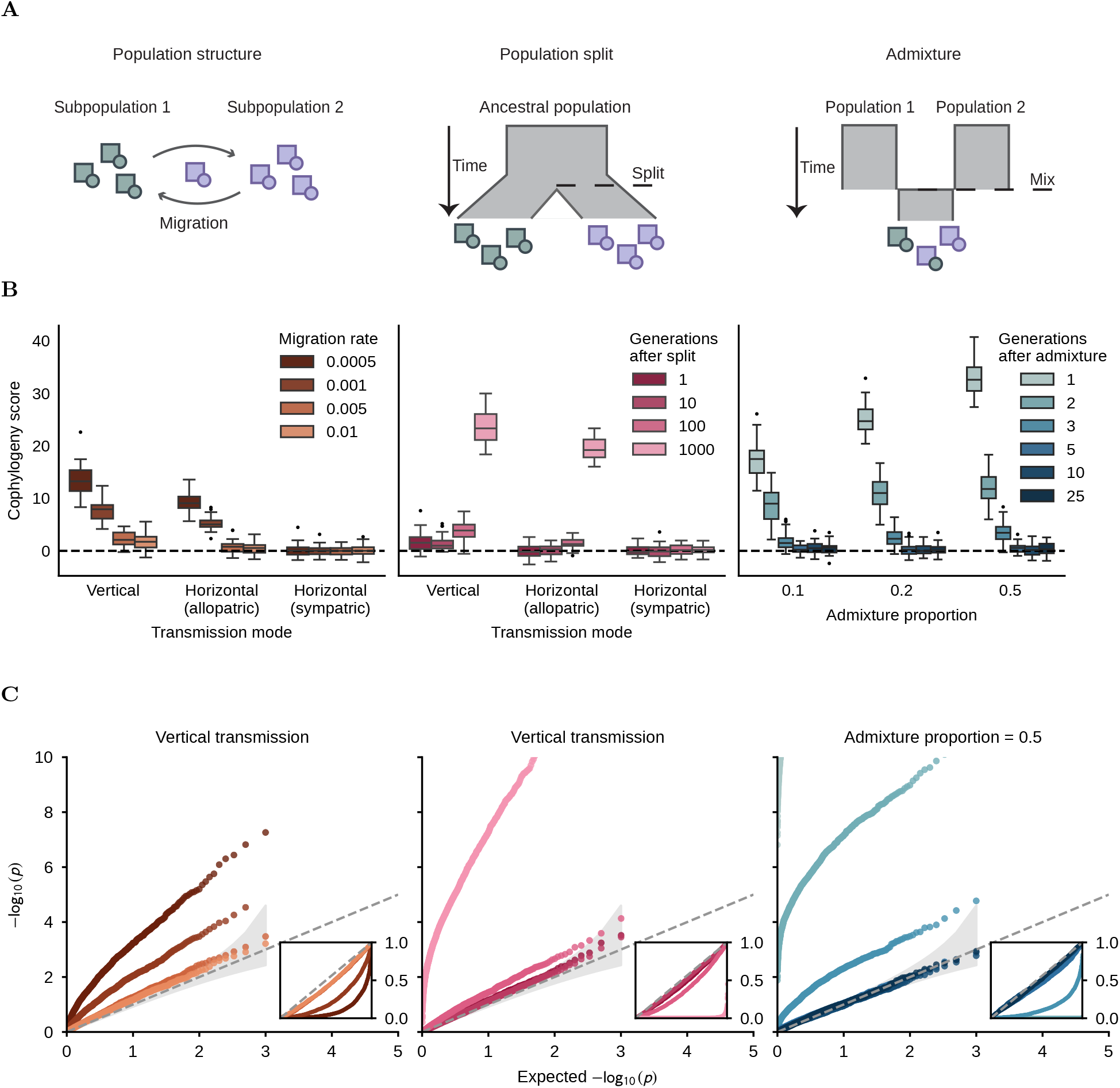
The effect of population structure on cophylogeny. A) Graphical depictions of our models of population structure, population split, and admixture. B) The effect of population structure on cophylogeny scores. Each boxplot represents the cophylogeny scores of 32 simulated replicates for a given set of parameters across varying demographic models and transmission modes. C) The effect of population structure on locus-specific cophylogeny. Q-Q plots for simulations of strict vertical transmission across demographic models. Cophylogenetic *p*-values were sampled at every 1 kilobase across simulated genomes and ranked by significance. Each point in the plot reflects the median ranked *p*-value across all 32 simulated replicates per parameter set. Inset Q-Q plots display the raw *p*-values. Shaded regions represent the 95% confidence intervals of expected *p*-values under the null hypothesis. Admixture Q-Q plot reflects an admixture proportion of 0.5, and cophylogenetic *p*-values for 1 and 2 generations after admixture largely fall outside of the plot limits.

In both models of population structure, we found that genome-wide cophylogeny only arose when transmission was vertical or horizontal allopatric (Figure 3B). When symbionts could freely move across subpopulations, as in our horizontal sympatric simulations, cophylogeny did not arise between hosts and symbionts. This shows that genome-wide cophylogeny can result from shared population structure between hosts and symbionts, which can arise from vertical transmission and population structure in the host, or from barriers to gene flow between host subpopulations that horizontally-transmitted symbionts cannot traverse. Furthermore, we find that the strength of cophylogeny correlates with the strength of population structure in the host. In the two subpopulation model, lower levels of gene flow correspond to higher genome-wide cophylogeny. In the population divergence model, genome-wide cophylogeny increases with sampling time following the population split. In all cases of elevated genome-wide cophylogeny, we observe inflation of locus-specific *p*-values across the host ARG (Figure 3C).

We then investigated the influence of historical population structure on cophylogeny by simulating two ancestral populations of equal size (*N* = 5, 000) without gene flow for 30 *∗ N* generations, followed by an admixture event resulting in a single admixed population (*N* = 10, 000) (Figure 3A). For these simulations, transmission of symbionts was strictly vertical, and we considered various proportions of ancestry contributed from Population 1 to the admixed population (0.1; 0.2; 0.5). At multiple generations following the admixture event (1; 2; 3; 5; 10; 25), we sampled 100 individuals and their symbionts, recapitated the host ARGs using `pyslim` with an ancestral effective population size of *N* = 10, 000 (Gopalan et al. 2025), and measured cophylogeny.

Across all admixture proportions, cophylogeny scores were highly elevated following the admixture event, but decreased rapidly within a few generations (Figure 3B). The proportion of admixture also affected the strength of cophylogeny scores, with higher admixture proportions resulting in stronger signals. We find similar trends for the inflation of locus-specific *p*-values (Figure 3C). Altogether, these results demonstrate how shared population structure between hosts and symbionts, both historical and contemporary, can drive strong cophylogeny scores and inflate locus-specific *p*-values.

To assess population structure within host populations, we computed *F_IS_* on the host samples from our simulations, which calculates the deviation from Hardy-Weinberg equilibrium (HWE) within a sample (Section 4.1). While *F_IS_*captures contemporary population structure, such as in our two subpopulation and population split models, it is not elevated in the admixture scenarios, where random mating following admixture immediately restores HWE (Figure S2).

### 2.4 Allelic incompatibility on standing variation

To investigate the effects of allelic incompatibility between specific genetic variants in hosts and symbionts, we added allelic states for host-symbiont pairs into our model. First, we simulated a panmictic population with constant population size (*N* = 10, 000), strict vertical transmission of symbionts, and a mutation rate of 10*^−^*^8^ in both hosts and symbionts. After 15 *∗ N* generations, we randomly selected a segregating site in the host population with an allele frequency in the range of 0.4 to 0.6, and labeled the derived allele “A” and the ancestral allele “B”. Next, we looked for mutations in the symbiont genomes within an allele frequency range of 0.2 to 0.8, and randomly selected a target mutation, labeling the derived allele “a” and the ancestral allele “b”. We then applied selection on the combined allele states of hosts and symbionts. In particular, we set matching homozygotes (AAa; BBb) and heterozygotes (ABa; ABb; BAa; BAb) as viable, and mismatched homozygotes (AAb; BBa) as lethal (Figure 4A). At the onset of selection and several generations after (0; 5; 10; 100), we sampled 100 individuals and their symbionts and measured cophylogeny. In each of the 32 simulated replicates, we found that allelic incompatibility drove a single pair of compatible alleles toward fixation, suggesting that host-symbiont allelic incompatibilities are transient within a panmictic population (Figure 4B). During the first few generations of selection, however, we observed a strong locus-specific cophylogenetic signal surrounding the focal locus in the host, which vanishes once genetic variation is lost (Figure 4C). These simulations demonstrate scenarios in which allelic incompatibilities drive locus-specific cophylogenetic signals in the host, and indicate the potential of locus-specific signals of cophylogeny to uncover genomic variation in the host that interacts with genetic variants in the symbiont.

**Figure 4:**
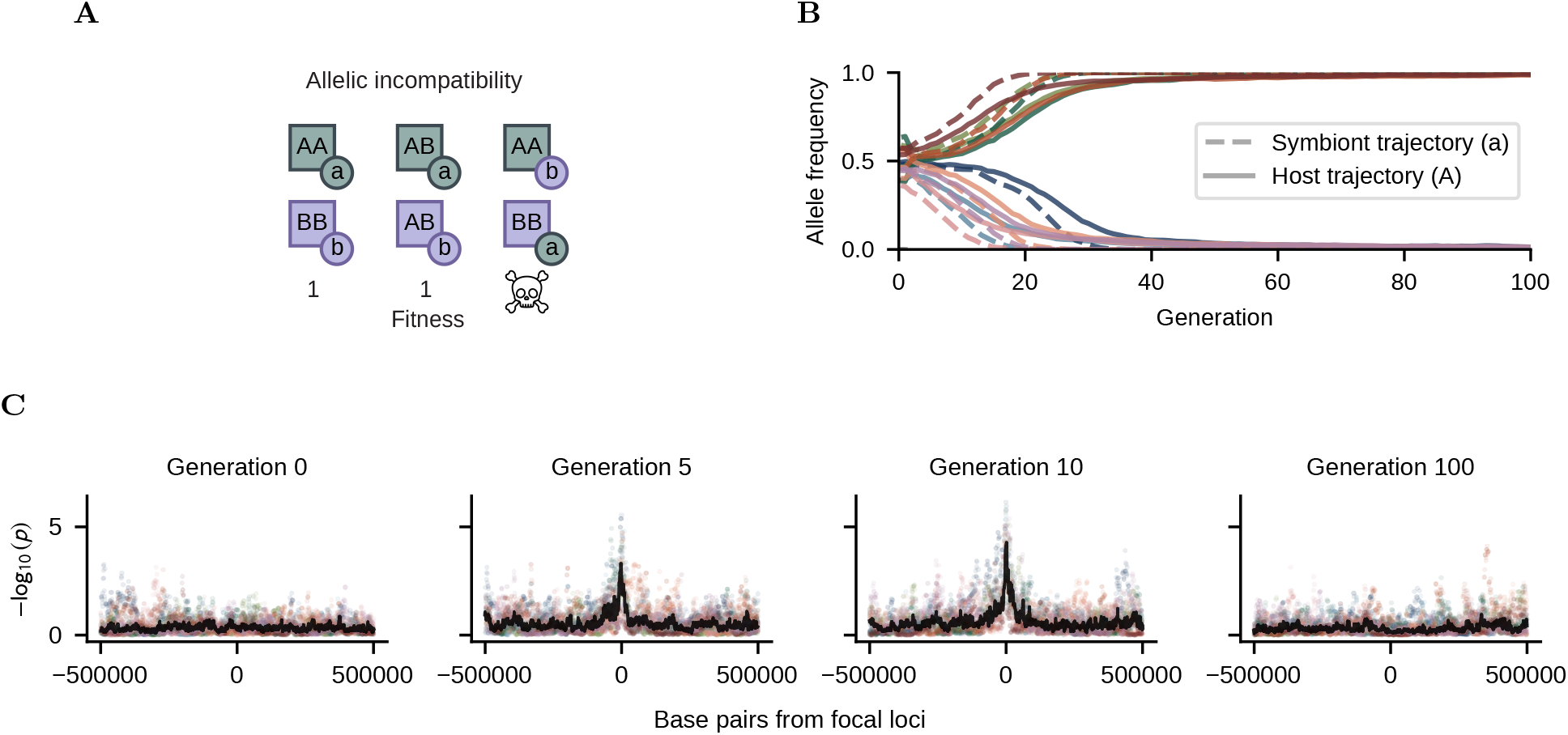
Allelic incompatibilities and locus-specific cophylogeny. A) Graphical representation of the allelic incompatibility model, where mismatching alleles between host and symbiont cause decreases in fitness. B) Allele frequency trajectories of host and symbiont alleles under selection for allelic incompatibility. Solid lines are allele frequency trajectories of host allele A and dashed lines reflect the allele frequency trajectories of symbiont allele a, colored by separate simulated replicate. C) Locus-specific cophylogenetic signals surrounding the focal locus in the host genome. Individual points are cophylogenetic *p*-values recorded every kilobase, colored based on simulated replicate, and median *−* log_10_(*p*) at each point across simulations is plotted as a black line.

### 2.5 Allelic incompatibility in admixed populations

Novel alleles in hosts and symbionts may arise and fix within diverged sub-populations, but cause incompatibilities when they are introduced into the same population by admixture. Such scenarios can give rise to genetic variation resembling Dobzhansky–Muller incompatibilities (Dobzhansky 1937; Muller 1942). To explore the influence of such incompatibilities on cophylogeny, we extended the admixture model from Section 2.3 to include allelic incompatibilities (Figure S3A). At a specific locus, hosts and symbionts within Population 1 were all set to have the derived allele (“A” in hosts and “a” in symbionts), and hosts and symbionts in Population 2 were all set to have the ancestral state (“B” in hosts and “b” in symbionts). Upon admixture, we set matching homozygotes (AAa; BBb) and heterozygotes (ABa; ABb; BAa; BAb) as viable, and mismatched homozygotes (AAb; BBa) as lethal (Figure S3A). We observed that cophylogeny scores persisted longer in simulations with an allelic incompatibility compared to simulations without, likely caused by the reinforcement of pre-admixture population boundaries (Figure S3B). Again, in each of the 32 simulated replicates, a single pair of compatible host-symbiont alleles was driven toward fixation (Figure S3C). Under varying proportions of admixture, we observed that alleles from the minor ancestry were typically lost, compatible with empirical observations of incompatibilities in admixed populations (Schumer et al. 2018). Locus-specific cophylogenetic signals at the focal locus of the host were not elevated compared to the genomic background (Figure S3D), demonstrating limitations in detecting locus-specific cophylogenetic signals when incompatibility alleles are strongly linked to diverged genomic backgrounds, for example, in recently admixed populations.

### 2.6 Mitonuclear cophylogeny in samples from the 1000 Genomes Project

To apply our approach to an empirical dataset of hosts and vertically transmitted symbionts, we obtained paired nuclear and mitochondrial genomes from human individuals in the 1000 Genomes Project (1KGP), a dataset of 2,504 genomic samples from 26 populations across the globe (Auton et al. 2015; Byrska-Bishop et al. 2022). We measured cophylogeny between the mitochondrial tree and the whole-genome ARG within each population in the 1KGP, inferred using `Relate` (Figure 5), see Section 4.2 for details. Of the 26 populations in the 1KGP, 13 displayed no genome-wide signal of cophylogeny (*p >* 0.05), in line with the results from our simulations that vertical transmission alone does not cause cophylogeny in large panmictic populations (Section 2.2). Only samples from five populations—Indian Telugu in the UK (ITU), Sri Lankan Tamil in the UK (STU), Punjabi in Lahore, Pakistan (PJL), African Ancestry in Southwest US (ASW), and Luhya in Webuye, Kenya (LWK)—displayed genome-wide cophylogeny with significance surpassing a multiple testing threshold (*p <* 2 *∗* 10*^−^*^3^, Bonferroni correction for 26 tested populations). Across all 1KGP populations, ITU, STU, and PJL displayed the three highest values of *F_IS_* (Figure S4), suggesting contemporary population structure or ongoing admixture within these groups. In contrast, ASW and LWK displayed comparatively low *F_IS_*(*<* 0.01) but strong cophylogeny scores, similar to our simulations of admixed or weakly structured populations (Section 2.3). Other admixed populations from the Americas (Zaidi and Makova 2019; Auton et al. 2015)—Mexican Ancestry in Los Angeles, California (MXL), Colombian in Medellin, Colombia (CLM), Puerto Rican in Puerto Rico (PUR), and African Caribbean in Barbados (ACB)—also displayed elevated cophylogeny scores.

**Figure 5:**
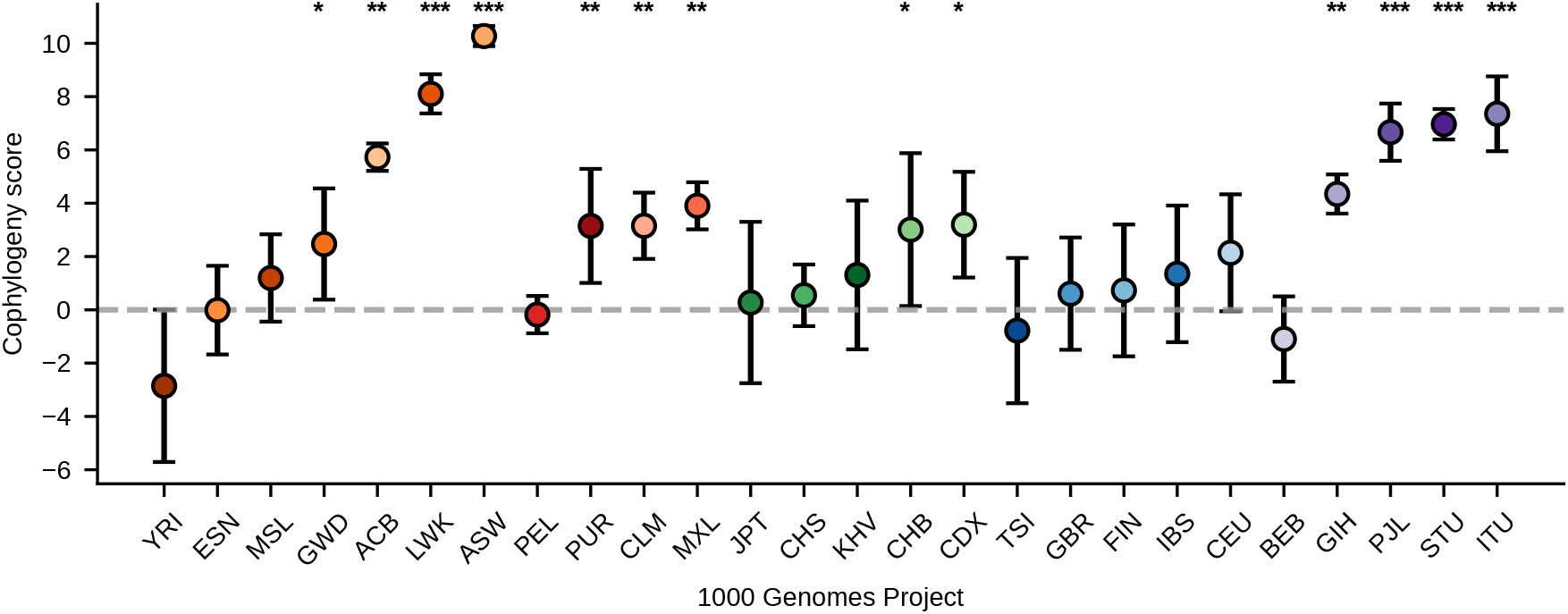
Mitonuclear cophylogeny scores across samples in the 1000 Genomes Project (1KGP). Each point represents the genome-wide cophylogeny score for each population in the 1KGP. Error bars represent 95% confidence intervals and were generated via block-jackknife (Busing et al. 1999) using chromosomes as blocks. Points are colored by population. Stars reflect *p*-values for significance: * *p <* 0.05, ** *p <* 0.01, *** *p <* 0.001.

To explore locus-specific signals of mitonuclear cophylogeny, we restricted our subsequent investigation to the 13 populations in the 1KGP without a significant genome-wide signal of mitonuclear cophylogeny (*p >* 0.05) (Figure 5). Using a Bonferroni correction, we set the population-specific genome-wide significance threshold to 0.05 divided by the total number of genealogies in each population’s inferred ARG, since each tree is tested for cophylogeny. Across the 13 populations analyzed, two genealogies displayed locus-specific cophylogenetic signal above their respective genome-wide significance thresholds. The strongest signal came from a genealogy in the ARG of Toscani in Italy (TSI) (*p* = 2.8 *∗* 10*^−^*^8^), which spanned 4,640 base pairs of intronic and exonic regions of *BCAR3*, a gene involved in antiestrogen resistance of human breast cancer cells (Agthoven et al. 1998). The second signal arose from a genealogy in the ARG of the Bengali in Bangladesh (BEB) (*p* = 5.9 *∗* 10*^−^*^8^), a 96-base-pair region within the intron of *CTNNA2*, which encodes *α*N-catenin primarily expressed in the brain (Schaffer et al. 2018). Neither signal maintained significance when accounting for multiple testing across all 13 populations (*p <* 4.8 *∗* 10*^−^*^9^).

To investigate mitonuclear interactions, we surveyed locus-specific cophylogenetic signals within nuclear genes involved in the synthesis of adenosine triphosphate via oxidative phosphorylation (OXPHOS), which relies on close coordination between nuclear and mitochondrial encoded proteins in humans (Zaidi and Makova 2019; Pfanner et al. 2019). For each of the 13 populations, we extracted the highest locus-specific cophylogenetic signal, and searched for overlap with one of 187 OXPHOS genes (*<* 0.16% of the genome, thus *p* = 0.02 for at least one overlap). In the sampled Japanese in Tokyo, Japan (JPT), the strongest cophylogenetic signal genome-wide was exhibited by a genealogy that directly overlapped *NDUFS7* (Figure 6A), a gene involved in complex I of the mitochondrial respiratory chain and implicated in mitochondrial diseases in humans (Lebon et al. 2007). To investigate this cophylogenetic signal in more detail, we employed the `MutualClusteringInfo` function from the `TreeDist` package (Smith 2020), which revealed that a shared clade between the mitochondrial tree (containing the D4, G2 and M9 haplogroups) and the focal genealogy within *NDUFS7* contributed most to the elevated cophylogenetic signal (Figure 7). In the focal genealogy, we observed 8 intronic segregating variants, and found that 4 displayed a linear relation to the presence or absence of the D4 haplogroup (linear regression, raw *p*-values *<* 6.3 *∗* 10*^−^*^3^), potentially indicative of a mitonuclear interaction in the population represented by the JTP sample. We extended our survey to include 100 kilobases upstream and downstream of OXPHOS genes, which revealed that the elevated signal around the strongest locus-specific cophylogenetic signal within the samples from the Iberian populations in Spain (IBS) covers the gene *BCS1L*, which encodes a protein involved in assembly of complex III of the mitochondrial respiratory chain and human mitochondrial disorders (Hikmat et al. 2021) (Figure 6A).

**Figure 6:**
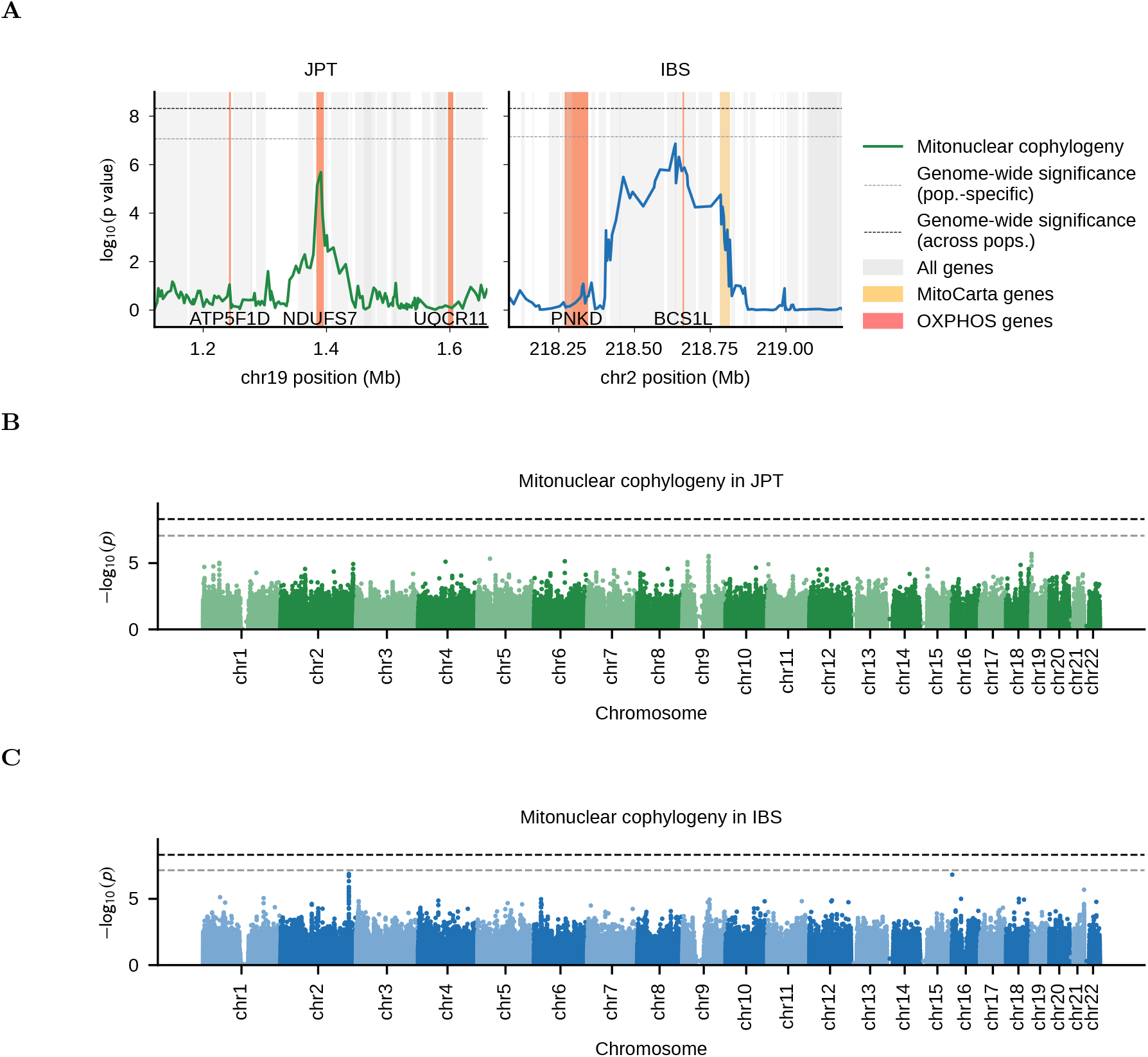
Locus-specific mitonuclear cophylogeny in JPT and IBS samples. A) Strongest locus-specific cophylogenetic signals across JPT and IBS genomes. Solid lines reflect the significance of cophylogeny across each respective genomic region. Genes (including coding and non-coding regions) are shaded in gray, genes involved in oxidative phosphorylation are shaded in red and labeled, and remaining MitoCarta genes are shaded in yellow. B) and C) Locus-specific cophylogeny for JPT and IBS genome-wide. *P* -values reflect the significance of cophylogeny between the mitochondrial tree and each genealogy in the host ARG. The x-axis shows the genomic position of the respective genealogy in the human genome.

**Figure 7:**
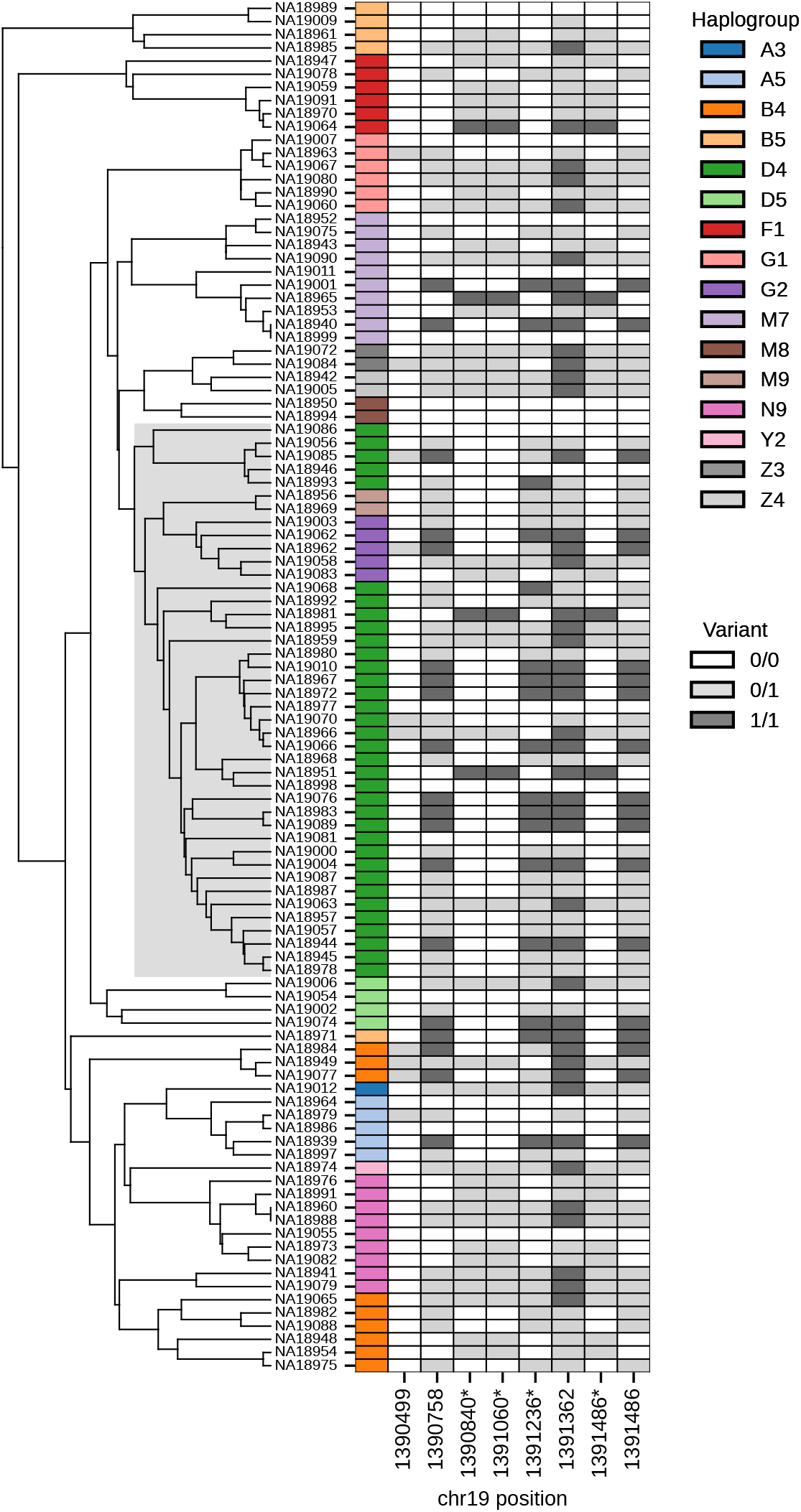
Cophylogeny between mitochondrial tree and focal genealogy in JPT sample. On the left is the UPGMA mitochondrial tree from the JPT (1KGP) samples. The grey box represents a shared clade between the mitochondrial tree and the focal genealogy that elevated the cophylogenetic signal (MutualClusteringInfo). Tip labels serve as the y-axis for the heatmap on the right, where the first column is colored by mitochondrial haplogroup, and the remaining eight columns display standing nuclear genetic variation within the genomic region covered by the respective focal genealogy of the ARG. Genomic variants are colored gray for heterozygotes, white for homozygous reference alleles, and dark gray for homozygous alternate alleles. Variants with a linear relation to the presence or absence of the D4 haplogroup are marked with a star (linear regression, raw *p*-values *<* 6.3 *∗* 10*^−^*^3^).

To test for enriched locus-specific cophylogenetic signals in functional genomic regions or classes of genes, we recorded the cophylogenetic signals at 1 kilobase intervals across the host genome, and extracted the loci with the top 1% strongest locus-specific signals within each population. We tested for enrichment (see Section 4.4) of protein coding genes (coding and non-coding regions), MitoCarta genes, and OXPHOS genes, as well as protein-coding exons, MitoCarta exons, and OXPHOS exons (Rath et al. 2021). Across all populations, we found no evidence of systematic enrichment, with only three significant signals of enrichment: exonic regions in CLM, protein-coding genes in IBS, MitoCarta genes in IBS; and one significant signal of depletion: OXPHOS genes for Esan in Nigeria (ESN) (Figure S5).

### 2.7 Mitonuclear cophylogeny in geographically structured samples

In Section 2.6, we measured mitonuclear cophylogeny in populations sampled from a single geographic location. To explore mitonuclear cophylogeny in geographically structured samples, we created sets of paired 1KGP populations and measured mitonuclear cophylogeny within sets. We selected ten populations (two from each 1KGP superpopulation) and created sets of all possible pairwise combinations, resulting in 45 sets of paired populations. Specifically, we selected from the superpopulation African Ancestry (AFR): Gambian in Western Division (GWD) and Yoruba in Ibadan, Nigeria (YRI); American Ancestry (AMR): CLM and PUR; East Asian Ancestry (EAS): Han Chinese South (CHS) and JPT; European Ancestry (EUR): IBS and TSI; South Asian Ancestry (SAS): Gujarati Indian in Houston, TX (GIH) and STU. With two geographically separated populations, we expected to find population structure in the hosts and corresponding mitonuclear cophylogeny (Section 2.3). Accordingly, we observed positive cophylogeny scores across all pairs, with a strong linear relation between population structure (assessed using *F_IS_*) and mitonuclear cophylogeny with ARGs built on chromosome 20 (*p*-value = 2.7 *∗* 10*^−^*^10^, *R*^2^ = 0.61) (Figure S6). Cophylogeny scores built on sets containing populations from the same superpopulation ranged from 1.7 (IBS and TSI) to 9.9 (GIH and STU), while sets containing populations from different superpopulations ranged in cophylogeny score from 17.3 (CHS and GIH) to 68.1 (YRI and TSI). Notably, both YRI and TSI displayed negative cophylogeny scores when analyzed individually (Figure 5), but a strong positive signal when analyzed together. These results reinforce the importance of population structure in cophylogeny scores, and demonstrate how geographic structure combined with vertical transmission can generate a range of genome-wide signals of cophylogeny.

## 3 Discussion

Here, we explored microevolutionary cophylogeny by considering variation in ancestral relations across the host genome. We do so by comparing a symbiont evolutionary tree to all local genealogies in the host ARG. Our simulations and empirical results demonstrate that vertical transmission alone does not generate cophylogeny between host ARGs and symbionts in large panmictic populations. Instead, we find that microevolutionary cophylogeny is primarily driven by shared population structure. Shared population structure can arise between hosts and symbionts when there is population structure in the host and vertical transmission of the symbionts, or when the barriers to gene flow in the hosts also prevent gene flow between symbionts, even if the symbionts are transmitted horizontally. These results suggest that microevolutionary cophylogeny between host ARGs and symbionts is not indicative of transmission mode within a single host species.

By measuring cophylogeny across all genealogies in the host ARG, we explored cophylogenetic variation across host nuclear genomic loci. Through simulations, we demonstrated that transient signals of locus-specific cophylogeny arise from allelic incompatibilities between host and symbiont genetic variants, particularly in unstructured populations with standing genetic variation. Thus far, there has been little evidence of mitonuclear incompatibilities in humans (Eyre-Walker 2017). However, most analyses search for incompatibilities between recently admixed populations (Rishishwar and Jordan 2017; Zaidi and Makova 2019; Zaidi et al. 2023). Our empirical analysis of mitochondrial and nuclear cophylogeny in putatively unstructured human populations illuminated a promising cophylogenetic signal in JPT between the mitochondrial tree and a nuclear encoded protein, *NDUFS7*, involved in complex I of the mitochondrial respiratory chain. Whether this signal reflects a mitonuclear incompatibility requires further analysis, and could benefit from the inclusion of samples with mitochondrial diseases, such as Leigh syndrome (Lebon et al. 2007). Our simulations and empirical results highlight a potential role of standing variation in mitonuclear incompatibilities within populations, especially if there are recent changes in environment and adaptive pressures.

Our approach is broadly applicable and can be used in many other host-symbiont systems, as long as the quality of the host genomic data allows for the reconstruction of ARGs. While mitochondria are exclusively transmitted maternally and can be well described with a single genealogical tree, other symbionts may experience extensive recombination, for example through horizontal gene transfer. In other host-symbiont systems, multiple genealogies for different genomic regions of the symbiont rather than a single tree might have to be considered to provide novel insights into the evolutionary mechanisms of host-symbiont genomic interactions.

## 4 Materials and Methods

### 4.1 Computing *F_IS_*

To measure *F_IS_*in the simulated data, we used `msprime` (Baumdicker et al. 2022) to simulate mutations across the host ARGs for each simulated replicate, with a mutation rate of 10*^−^*^7^. We then exported the mutation information to a vcf-file, filtered for biallelic single nucleotide polymorphisms, and calculated *F_IS_* for each simulation replicate using `plink2` (Chang et al. 2015) and a custom python script, following Hartl and Clark (2007, Ch. 4).

To measure *F_IS_* on 1KGP data, we filtered for single nucleotide polymorphisms with minor allele frequency *>* 0.01, and extracted sites passing an accessibility mask (see Section 5). We then used plink2 and a custom python script to measure *F_IS_*, again following Hartl and Clark (2007, Ch. 4).

### 4.2 Inferring ancestral recombination graphs for the 1000 Genomes Project

We obtained the whole-genome sequencing data for the 1KGP, resequenced by the New York Genome Center (Byrska-Bishop et al. 2022). We then used `Relate` (v1.2.4) to infer ARGs for each population individually and select population pairs, using the recommended parameters, recombination map, mutation rate, accessibility mask, and coalescence rates for GRCh38 (Speidel et al. 2019). For the mitochondrial tree, we computed distances between mitochondrial genomes (included in the 1KGP data) using the Kimura two-parameter distance (Kimura 1980), and built the genealogical tree using unweighted pair group method with arithmetic mean (UPGMA; Felsenstein 2004, Ch. 11).

### 4.3 Measuring cophylogeny between a symbiont tree and a host ARG

To measure distances between phylogenetic trees, we used the function `TreeDistance` (Smith 2020), which quantifies topological distance between two trees by comparing shared bipartitions. This metric requires identical leaf sets between the two trees. However, haploid symbiont trees have a single leaf per individual (hosting the respective symbiont), whereas ancestral recombination graphs treat diploid chromosomes independently, resulting in two leaves per individual. Thus, before comparing symbiont trees and ARGs, we converted each chromosome-based tree in the ARG to an individual-based tree. To do this, we treated each individual as a set of two leaves, containing both their maternal and paternal chromosomes, and computed genetic distances between individuals by averaging the values of the time to the most recent common ancestor (*T*_MRCA_) between the respective sets. We calculated distances between all individuals and applied UPGMA to generate individual-based trees.

To obtain genome-wide cophylogeny scores, we assessed the strength of cophylogeny between a symbiont tree and the full host ARG through permutation testing. We permuted the leaves of the symbiont tree 1000 times. For each of the *n* local genealogical trees in the ARG, we measured `TreeDistance` *d_i,j_* between the *i*-th tree in the ARG and the *j*-th permuted symbiont tree, as well as `TreeDistance` 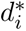 between the *i*-th tree in the ARG and the true symbiont tree. Genome-wide cophylogeny, “cophylogeny score”, was then defined as follows:

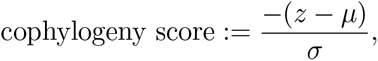

where the sign in the numerator is chosen such that a greater “distance” leads to a lower “cophy-logeny” score, 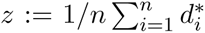 is the mean `TreeDistance` between the ARG trees and the true symbiont tree, 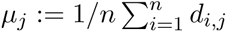 is the mean `TreeDistance` between the ARG trees and the *j*-th permuted tree, 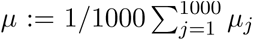 is the average of these means across permutations, and *σ* is the respective standard deviation. We assessed the significance of the genome-wide cophylogeny score using a *p*-value computed as the empirical percentile of the true mean value *z* among the permuted mean values *µ_j_*.

Permutation testing can also be used to assess the significance of cophylogenetic signals for a specific local genealogical tree in the ARG. However, obtaining reliable *p*-values that can be corrected for multiple testing among many local genealogies would require a prohibitive amount of permutations. We thus fit a parametrized distribution to a smaller set of permuted values to approximate the null distribution under permutation testing, a similar strategy as employed by Knijnenburg et al. (2009). To this end, we compared a given local genealogy in the ARG to 1000 symbiont trees with the leaves permuted to generate a null distribution of `TreeDistance` scores specific to this local genealogy. We then fit a normal-inverse Gaussian (NIG) to the set of permuted distances, and calculated a *p*-value for the true distance based on the fit distribution. To demonstrate that this approach approximates permutation testing well, we simulated 64 pairs of marginal coalescent trees for a population of size *N* = 10, 000 using msprime (Baumdicker et al. 2022), for several different sample sizes (50; 100; 200). For each pair, we applied 100,000 permutations to the leaves of one of the two trees, and computed `TreeDistance` for each permuted pair to obtain a null distribution of `TreeDistance` values under permutation and reported the empirical *p*-value. Further, for each pair, we subsampled 1000 of the permutations, fit a NIG distribution to this subsample, and computed approximate *p*-values for all 100,000 permuted `TreeDistance` values using this fit distribution. The *p*-values obtained using the NIG distribution closely reflected the *p*-values obtained from direct permutation testing (Figure S1).

### 4.4 Enrichment

To assess enrichment of genomic elements, we recorded locus-specific cophylogenetic *p*-values every 1kb along the genome within each population. We then extracted the top 1% of loci with the strongest cophylogenetic signals within each population and measured the fraction of genomic elements within the top loci. We computed enrichment by dividing this fraction by the fraction of genomic elements found across all loci where *p*-values were recorded. We used GENCODE v49 to obtain genomic features and gene annotations (Mudge et al. 2025), and MitoCarta3.0 to label nuclear encoded mitochondrial proteins, as well as proteins involved in OXPHOS (Rath et al. 2021). We measured enrichment of genes (coding and non-coding regions) and exons for all protein-coding genes. We also measured enrichment of MitoCarta genes, and OXPHOS genes, and MitoCarta exons, and OXPHOS exons.

To assess significance of these enrichment values, we require an appropriate null distribution. Due to chromosomal linkage, locus-specific *p*-values at neighboring loci are correlated, and we thus need to reflect this correlation in the null distribution. First, to compute this correlation on a natural scale, we converted the locus-specific *p*-values to *z*-scores (corresponding quantiles of a standard normal distribution) and measured the autocorrelation between these *z*-scores at neighboring loci within each population. For each population, we then generated random sequences of correlated values sampled from a conditional bivariate normal distribution with the same autocorrelation, where the length of each sequence is equal to the total number of recorded loci across the genome. We considered these sequences our null genomes, since they maintain the correct correlation structure among the locus-specific values, but are otherwise ignorant of genomic features. For each population and each enrichment test, we generated 10,000 null genomes and measured enrichment in the top 1% of values each, providing a set of null enrichment values based on the correct underlying correlation structure within each population. We then used the rank of the true enrichment values within this null distribution to obtain a *p*-value for the observed enrichment.

## 5 Code and data availability

The code used for the data analyses and for generating the figures presented in this manuscript is available at https://github.com/steinrue/mitonuclear_cophylogeny (Release: “bioRxiv release v1.0.0”). Resulting locus-specific mitonuclear cophylogenetic *p*-values for samples from the 1000 Genomes Project (1KGP) are also available at https://doi.org/10.5281/zenodo.22679262. We downloaded phased genomes for the 1KGP from the International Genome Sample Resource (IGSR): https://www.internationalgenome.org/data-portal/data-collection/ `1000genomes_30x`. To download accessibility masks, recombination maps, ancestral sequences, and coalescent rates, we followed the instructions outlined at https://myersgroup.github.io/relate/, which includes accessibility masks available from the IGSR: https://www.internationalgenome.org/announcements/genome-accessibility-masks/, ancestral sequences from Ensembl release 86 (Dyer et al. 2024), and recombination maps adapted from https://bochet.gcc.biostat.washington.edu/beagle/genetic_maps/ (Browning et al. 2021).

## Acknowledgements

We want to thank Adam Fine for helpful comments and feedback on the manuscript. Computational resources were provided by the Center for Research Informatics, which is funded by the Biological Sciences Division at the University of Chicago with additional funding provided by the Institute for Translational Medicine, grant number UL1TR002389 from the National Institutes of Health. This work was supported in part by the National Institute of General Medical Sciences (NIGMS) of the National Institutes of Health under award R01GM146051 (MS, RH), award T32GM139782 (RH) and by grants from the NSF (DMS-2235451) and Simons Foundation (MPS-NITMB-00005320) to the NSF-Simons National Institute for Theory and Mathematics in Biology (NITMB).

## 7 Supplementary figures

**Figure S1:**
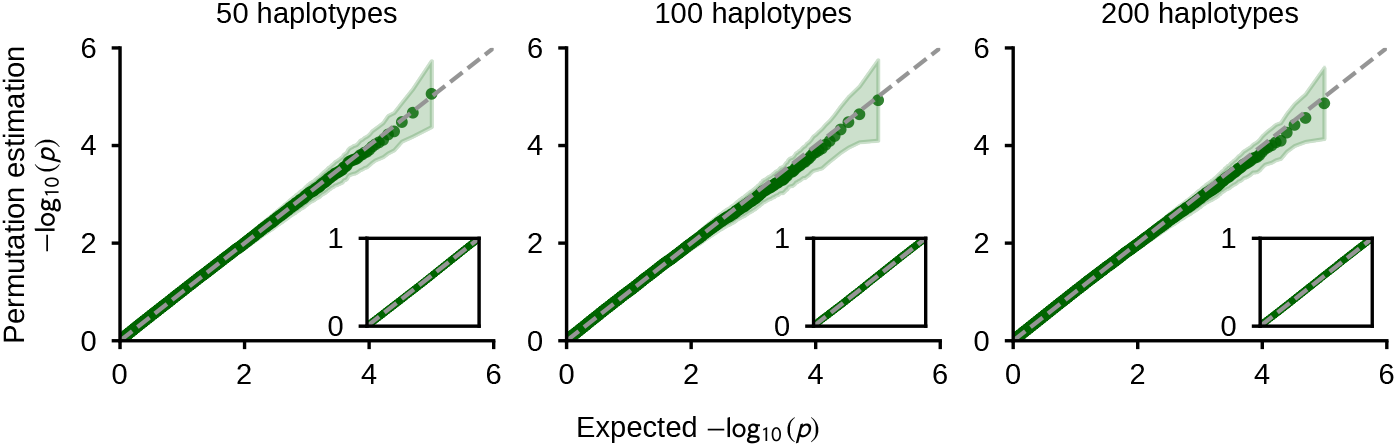
Validation of approximate permutation test. Empirical *p*-values for cophylogeny obtained from 100,000 permutations against *p*-values obtained from an NIG-distribution fitted to a subsample of 1,000 permutations. Each point reflects the median ranked *p*-value across 64 replicates. Shaded regions represent the standard deviation at each rank. Inset shows the same plot for raw *p*-values. Different plots for different number of leaves (sampled haplotypes) in the tree comparisons (50; 100; 200).

**Figure S2:**
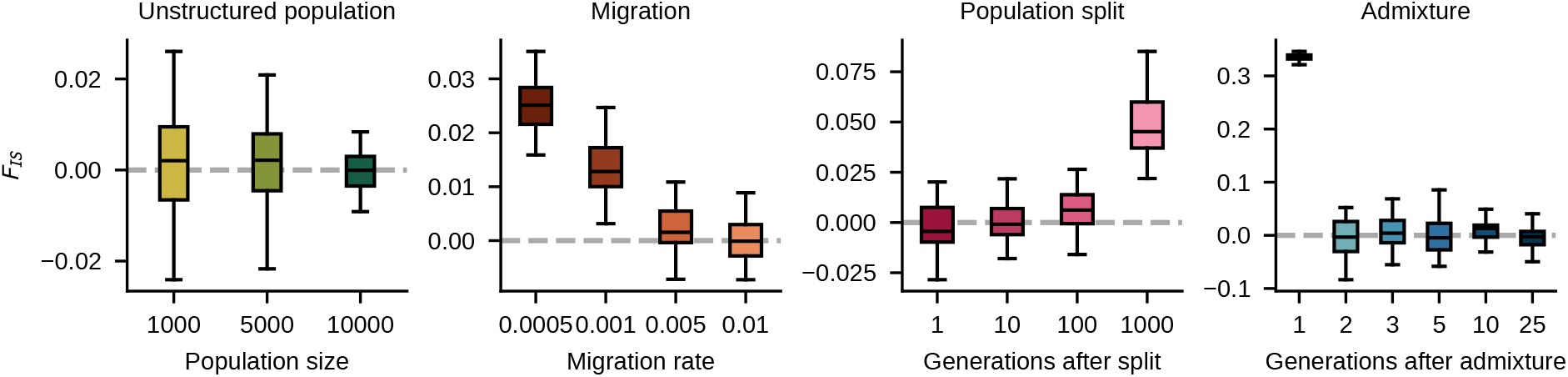
Assessing population structure from simulated host genomic data. Each boxplot shows the distribution of *F_IS_* values for the host samples calculated from 32 simulated replicates for a given set of parameters.

**Figure S3:**
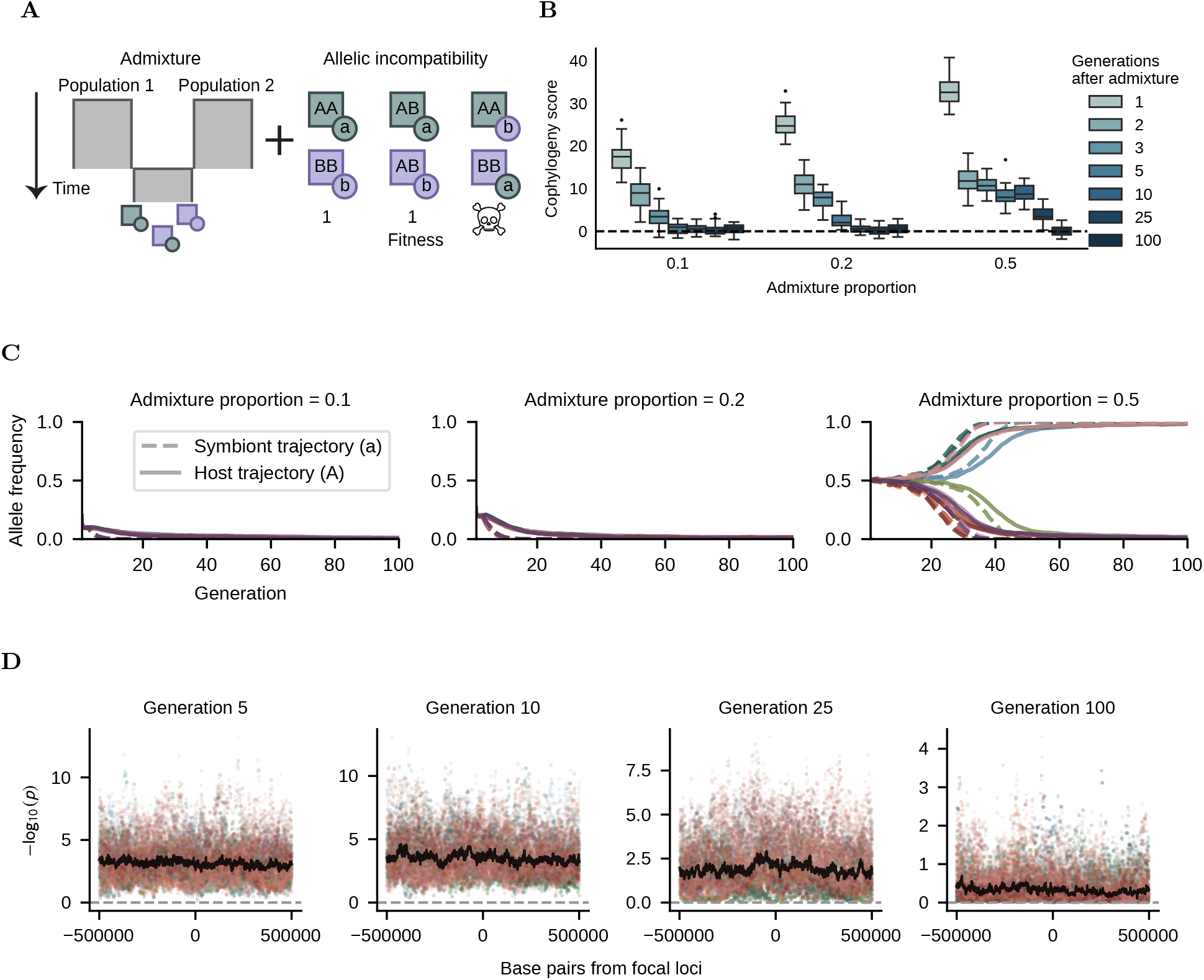
Admixture with an allelic incompatibility. A) Graphical representation of the admixture and allelic incompatibility model. B) The effect of admixture and allelic incompatibilities on cophylogeny scores. Boxplots of cophylogeny scores for 32 simulated replicates for a given set of parameters. C) Allele frequency trajectories of introgressed host and symbiont alleles from Population 1. Solid lines are allele frequency trajectories of host allele A and dashed lines reflect the allele frequency trajectories of symbiont allele a, colored by simulated replicate. D) Locus-specific cophylogenetic signals surrounding the focal locus in the host genome. Points are cophylogenetic *p*-values recorded every kilobase. Points are colored based on simulated replicate, and median *−* log_10_(*p*) at each point across simulated replicates is plotted as a black line. In contrast to Figure 4C, the focal locus does not show elevation in cophylogenetic signal compared to the background.

**Figure S4:**
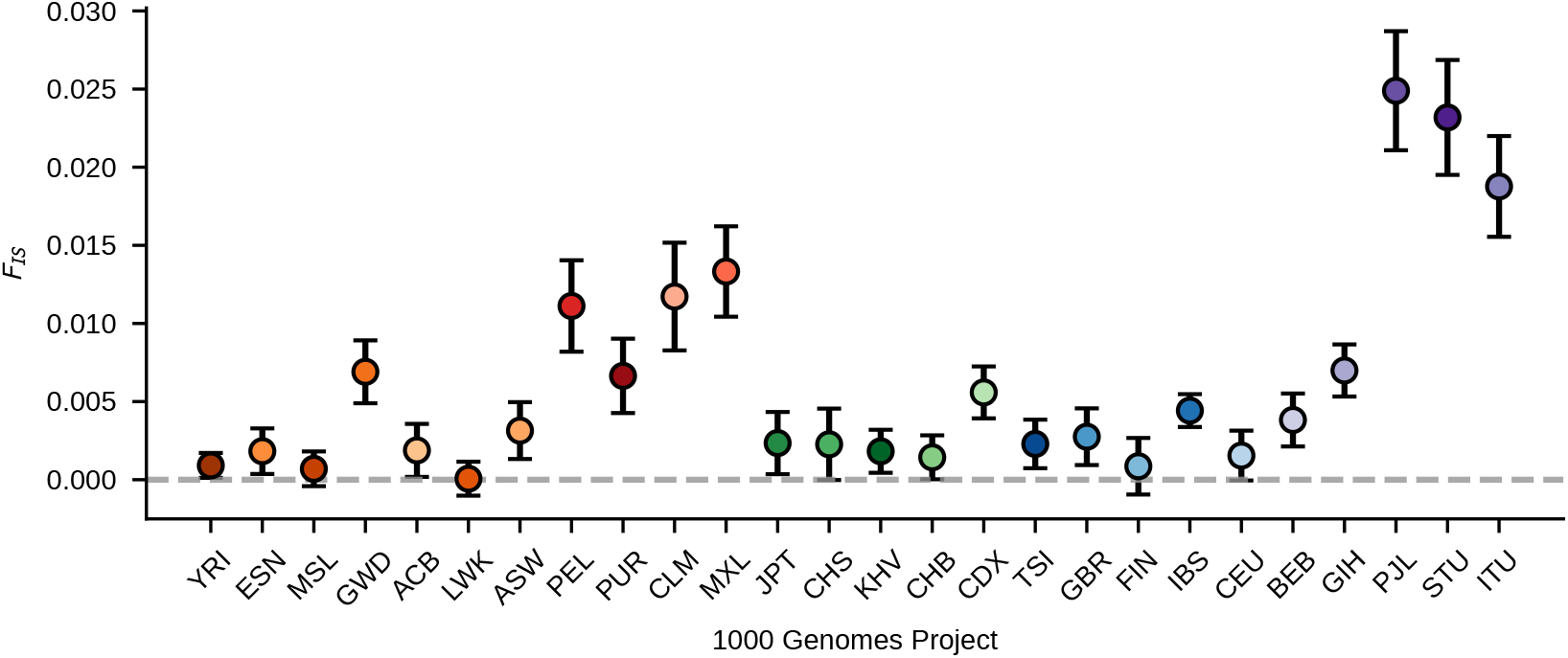
Assessment of population structure in samples from the 1000 Genomes Project (1KGP) using *F_IS_*. Each point shows *F_IS_*computed from the corresponding population sample in 1KGP data. Error bars represent 95% confidence intervals and were generated via block-jackknife (Busing et al. 1999) using chromosomes as blocks.

**Figure S5:**
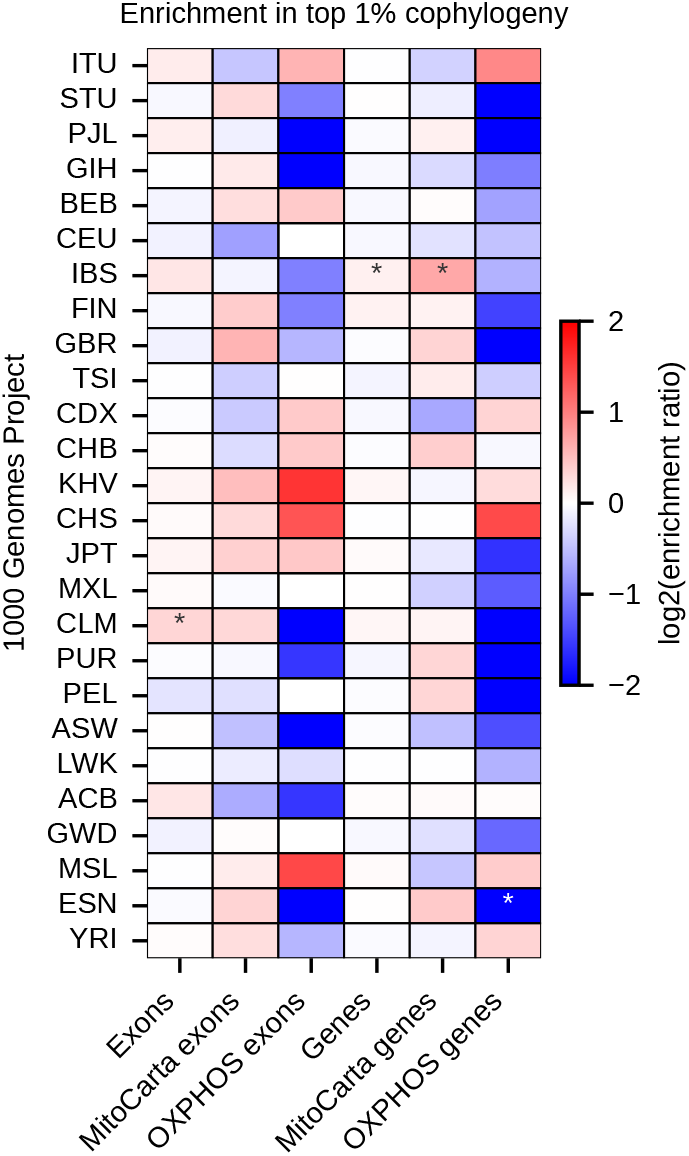
Functional enrichment of locus-specific cophylogenetic signals across human populations. Enrichment of functional classes in the top 1% of locus-specific mitonuclear cophylogenetic signals within each population. “Genes” refers to coding region + non-coding regions. Stars reflect significant *p*-values (* *p <* 0.05), Bonferroni-corrected for 26 * 6 tests.

**Figure S6:**
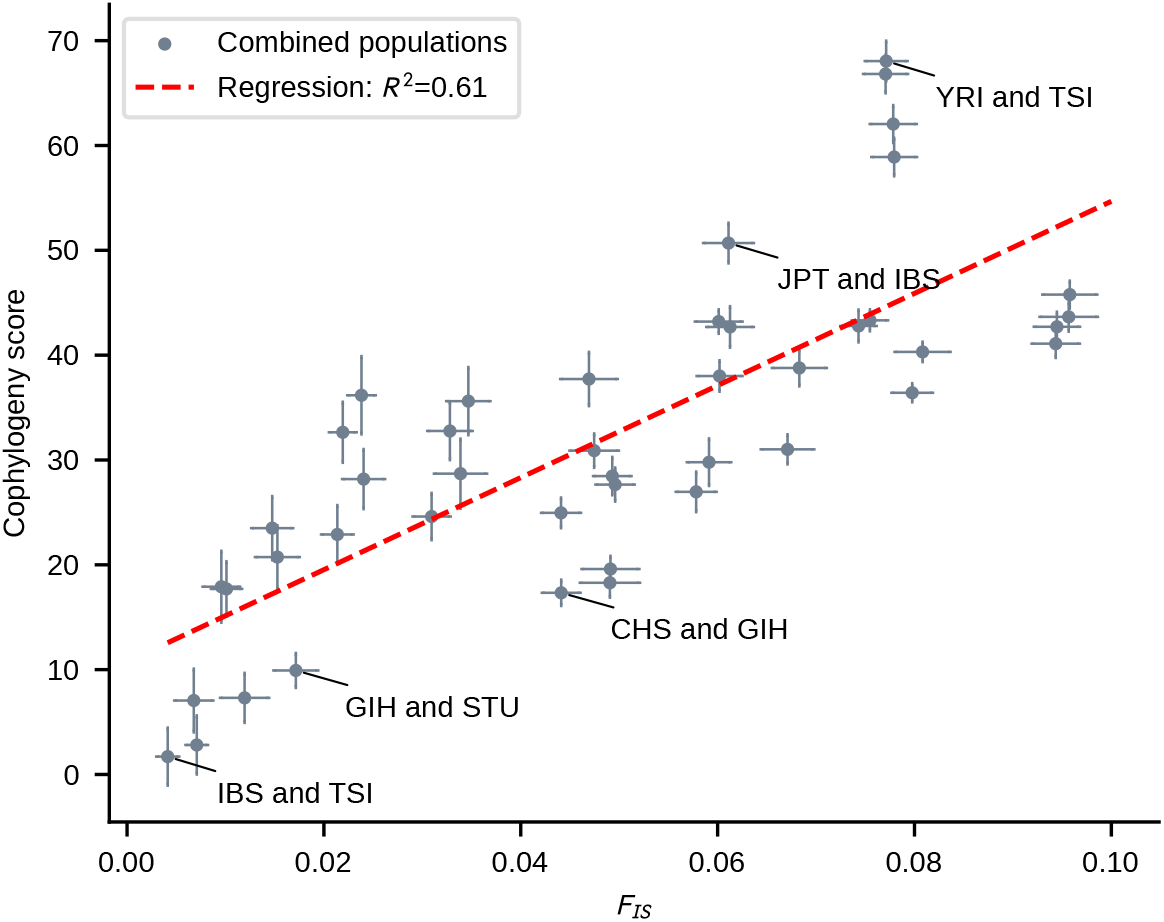
Cophylogeny scores in geographically structured samples. Each point reflects the cophylogeny score between an ARG built on chromosome 20 and the mitochondrial tree for a paired set of population samples, 5 of which are labeled. The x-axis shows the corresponding value of *F_IS_*, computed using the full host genome. Cophylogeny score error bars represent 95% confidence intervals and were generated through a block-jackknife approach (Busing et al. 1999) on chromosome 20 using genomic regions as blocks of size *≈* 3 Mb. *F_IS_* error bars represent 95% confidence intervals and were generated through a block-jackknife using chromosomes as blocks. A linear regression is shown as a red dashed line.

